# Size-dependent temporal decoupling of morphogenesis and transcriptional programs in gastruloids

**DOI:** 10.1101/2024.12.23.630037

**Authors:** Isma Bennabi, Pauline Hansen, Melody Merle, Judith Pineau, Lucille Lopez-Delisle, Dominique Kolly, Denis Duboule, Alexandre Mayran, Thomas Gregor

**Affiliations:** Department of Developmental and Stem Cell Biology, CNRS UMR3738 Paris Cité, Institut Pasteur, Paris, France; School of Life Sciences, Ecole Polytechnique Fédérale de Lausanne (EPFL), 1015 Lausanne, Switzerland; Center for Interdisciplinary Research in Biology (CIRB), Collège de France, CNRS, INSERM, Université PSL, Paris, France; Lewis-Sigler Institute for Integrative Genomics, Princeton University, Princeton, NJ, USA; Joseph Henry Laboratories of Physics, Princeton University, Princeton, NJ, USA

**Author notes:** equal contribution, ranked alphabetically (Isma Bennabi, Pauline Hansen). senior author (Denis Duboule, Alexandre Mayran, Thomas Gregor).

## Abstract

Understanding the interplay between cell fate specification and morphogenetic changes remains a central challenge in developmental biology. Gastruloids, self-organizing stem cell-based models of post-implantation mammalian development, provide a powerful platform to address this question. Here, we show that physical parameters, particularly system size, critically influence the timing and outcomes of morphogenetic processes. Larger gastruloids exhibit delayed symmetry breaking, increased multipolarity, and prolonged axial elongation, with morphogenesis driven by system size. Despite these variations, transcriptional programs and cell fate composition remain remarkably stable across a broad size range. Notably, extreme sizes show distinct transcriptional modules and clear shifts in gene expression patterns. Intriguingly, size perturbation experiments rescued the morphogenetic and pattern phenotypes observed in extreme sizes, demonstrating the remarkable adaptability of gastruloids to their effective system size. These findings establish gastruloids as versatile models for studying spatiotemporal dynamics in mammalian embryogenesis and reveal how physical constraints decouple transcriptional from morphogenetic programs.

## Introduction

Embryogenesis is a highly coordinated process orchestrating multiple lineage decisions together with morphogenetic changes. During gastrulation, vertebrates converge on a conserved body plan, a phenomenon described as the hourglass model [1, 2]. This critical event establishes the three germ layers (ectoderm, mesoderm, and endoderm) and major body axes [3]. Gastrulation relies on the interplay of gene expression, biochemical signals, mechanical forces, and geometry, precisely coordinated across spatial and temporal scales [4]. However, understanding how these factors are integrated to ensure robust embryonic development remains a longstanding challenge in developmental biology.

Stem cell-derived embryo models, such as gastruloids, have emerged as powerful tools to probe these mechanisms [5, 6, 7]. Gastruloids recapitulate key events of mammalian gastrulation, including germ layer specification and anteroposterior axis elongation. Developing without extraembryonic tissues, these self-organizing models offer unmatched experimental accessibility compared to native embryos [8, 9, 10, 11, 12, 13, 14]. A hallmark of gastruloid development is their robust ability to break symmetry and elongate along an anteroposterior axis, forming a posterior pole characterized by Brachyury expression. The conserved ability of gastruloids to self-organize and elongate an axis has been demonstrated across species, including human pluripotent stem cells [6, 15]and dissociated zebrafish embryo explants [16]. Notably, gastruloid axial elongation is a well-defined and highly reproducible morphogenetic process, positioning gastruloids as an ideal minimal system for quantitative developmental studies [17, 18, 19, 11, 14, 13]. Moreover, their conserved nature provides a unique platform to experimentally test the hourglass model, shedding light on how conserved morphologies can emerge through self-organization.

Another significant advantage of gastruloids is their scalability and amenability to environmental and physical perturbations. By adjusting the initial cell seeding number, gastruloids can be generated across a range of sizes, facilitating studies on how physical parameters, such as system size, modulate developmental processes. Early studies identified an optimal cell seeding number for symmetry breaking, Brachyury polarization, and subsequent anteroposterior elongation (Van den Brink et al., 2014). Additionally, size-dependent relative expansion of a SOX2-positive inner core of cells has been observed prior to axial polarization [20]. Conversely, studies has demonstrated that early Brachyury polarization and antero-posterior patterning are robust to size variations within a certain range, and several developmental genes main tain proportional expression patterns that scale with gastruloid length at later stages [21, 11]. Yet, whether size, cell fate, and morphogenesis are coupled remains unclear. Specifically, how do changes in system size influence the interplay between cell differentiation and morphogenesis? In this study, we address this question by systematically varying gastruloid size through changes in cell numbers and employing live imaging and transcriptional profiling techniques to monitor the temporal dynamics of these processes. Our results reveal that gastruloid size significantly impacts morphogenesis dynamics, driving changes in the timing of symmetry breaking, multipolarity, and axial elongation. Interestingly, despite pronounced morphogenetic changes, transcriptional states and cell fate composition remain largely unaffected within a broad size range (up to six-fold variation). However, at extreme sizes, we observe metabolic shifts and changes in gene expression patterns. Finally, we demonstrate that these changes are primarily governed by the effective size of the gastruloid rather than the initial cell seeding number.

This study highlights that, while transcriptional programs can be temporally decoupled from morphogenesis, the latter is strongly influenced by physical size constraints. Together, our findings establish gastruloids as an ideal experimental system for exploring the physical principles that govern the spatiotemporal dynamics of mammalian development.

## Results

### Gastruloid morphogenesis timing depends on the initial cell number

To investigate how size variation impacts gastruloid development, we generated gastruloids of various sizes by adjusting the initial number of seed cells, *N*_0_ (Fig. S1A). Mouse embryonic stem cells (mESCs) were cultured in 2i + LIF medium before seeding, ensuring a homogeneous cellular state [22] and highly reproducible gastruloid formation [20, 13, 11]. Following a WNT pathway activation pulse (Chiron), gastruloids seeded at the canonical size (*N*_0_ = 300) consistently break spherical symmetry, elongate along the anteriorposterior (AP) axis with high reproducibility, and eventually collapse [8, 9, 10]. We first characterized the size range that supports typical gastruloid development under our culture conditions. To do so, we tested a large range of initial cell numbers, spanning a 1200-fold range, from 25 to 30000 cells (Supplementary Movie 1). Smaller gastruloids (*N*_0_ ≤ 100 cells) elongate as early as 96 h but collapse by 144 h (Fig. 1A, Fig. S1B). Larger gastruloids (*N*_0_ ≥ ≥600) initially form multipolar structures and require more time to achieve uniaxial elongation, which rarely occurs at extreme sizes (Fig. 1A, Fig. S1B).

**Figure 1:**
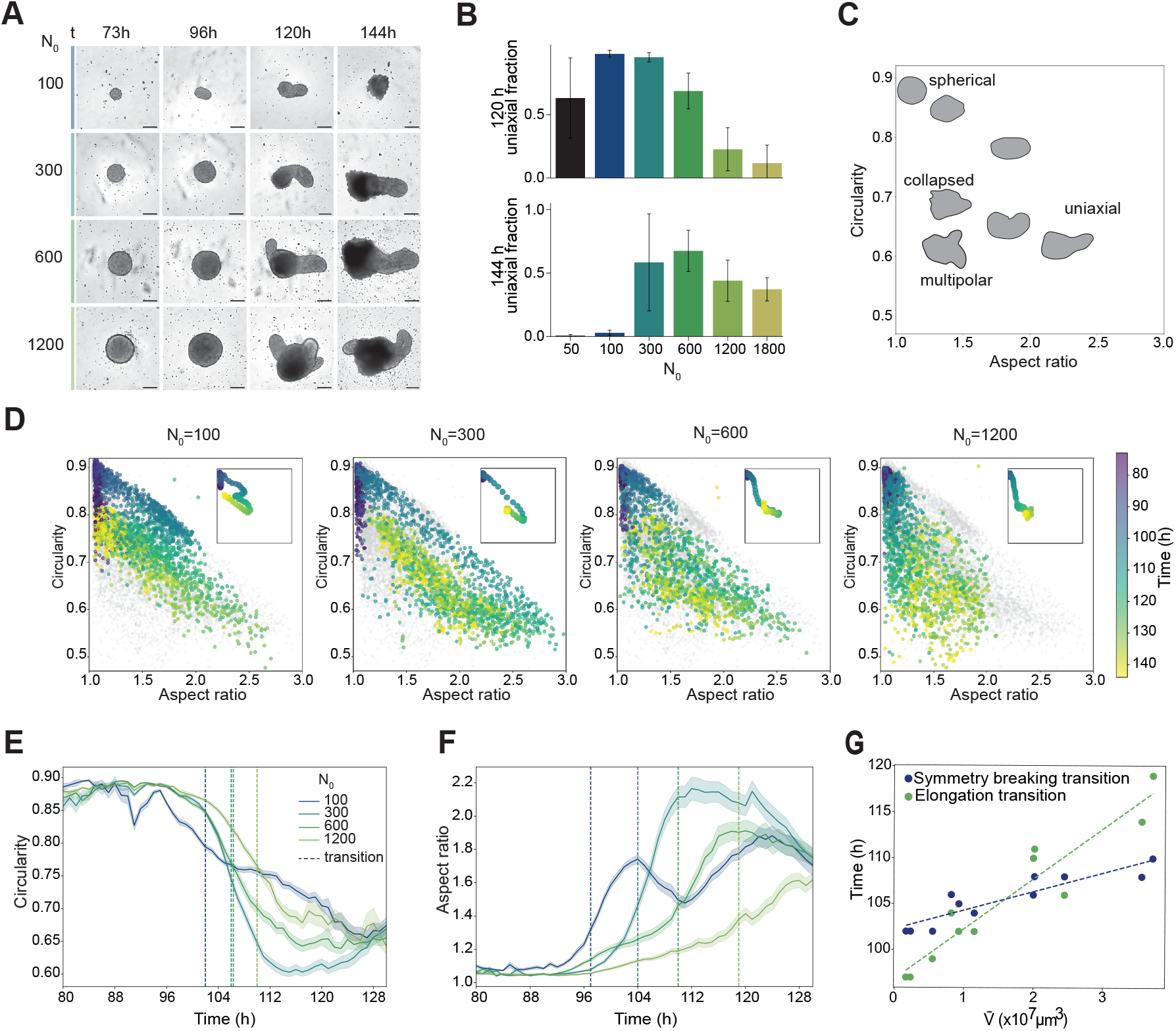
Gastruloid size governs morphogenesis timing. (A) Brightfield images showing gastruloid morphology at 72, 96, 120, and 144 h post-seeding across various initial cell numbers (*N*_0_). Scale bars = 200 *µm*. (B) Fraction of uniaxial gastruloids at 120 and 144 h for each *N*_0_. Bars represent mean *±* s.d. from three independent replicates. (C) Relationship between circularity and aspect ratio in different gastruloid morphologies: spherical, uniaxial (single elongation axis), multipolar (multiple axes), or collapsed (loss of tissue integrity). Example cartoon shapes extracted from real gastruloids. (D) Scatter plots of circularity versus aspect ratio for gastruloids with varying *N*_0_. Points represent individual gastruloids, colored by time (72–144 h). Insets show average morphological trajectories. Sample sizes: *N*_0_ = 100 (n = 41), *N*_0_ = 300 (n = 41), *N*_0_ = 600 (n = 36), *N*_0_ = 1200 (n = 40). (E, F) Temporal dynamics of circularity and aspect ratio (mean *±* s.e.m.) across *N*_0_ conditions. Dashed lines mark symmetry breaking and elongation transitions, determined via optimal partitioning (Fig. S1G and see Methods). Symmetry breaking, as evaluated from circularity, occurs at 102 h, 106 h, 106 h, and 110h for *N*_0_ = 100, 300, 600, and 1200, respectively. Elongation, as evaluated from aspect ratio, occurs at 97 h, 104 h, 110 h, and 119 h for *N*_0_ = 100, 300, 600, and 1200, respectively. (G) Transition times for symmetry breaking (blue) and elongation (green) as a function of gastruloid size at 73 h. Dashed lines indicate linear fits (*R*^2^ = 0.858 for symmetry breaking; *R*^2^ = 0.904 for elongation). See Supplementary Tables for sample sizes.

High-throughput live-imaging of gastruloids (*N*_0_ = 50 to 1800, 72 to 144 h post-seeding, three experimental batches) reveals that uniaxial elongation is most robust and reproducible for *N*_0_ = 100–300 cells (*>* 95%) (Fig. 1B). In contrast, larger gastruloids (*N*_0_ ≥ 600) initialized elongation along multiple axis, with a small fraction achieving uniaxial elongation by 120 h (Fig. 1A) and an increased fraction by 144 h (Fig. 1B). These findings demonstrate that while morphogenesis is supported across a wide range of sizes, the timing, robustness, and reproducibility of axial elongation are strongly size-dependent.

To quantify morphogenesis dynamics, we developed an automated segmentation method to analyze gastruloid shapes in live brightfield movies (Fig. S1C, see Methods). Circularity (a proxy for symmetry breaking) and aspect ratio (reflecting uniaxial elongation) were used as shape descriptors, effectively capturing gastruloid morphologies (Fig. 1C, Fig. S1C). At 72 h, gastruloids across all sizes are spherical, with circularity and aspect ratio values close to one (Fig. 1D). Smaller gastruloids (*N*_0_ ≤ 300) follow a trajectory with increasing in aspect ratio over time, indicating uniaxial elongation, until the trend reverses as the gastruloids begin to collapse. Larger gastruloids, however, exhibit higher multipolarity and reduced elongation, with lower circularity and aspect ratio values (Fig. 1D).

The dynamics of transition times show that morphological symmetry is maintained longer in larger gastruloids, as evidenced by delayed circularity reduction (Fig. 1E, Fig. S1D-E). Similarly, axial elongation is delayed by nearly a day in larger gastruloids (Fig. 1F, Fig. S1D-E-F).

From this analysis, we conclude that gastruloid size is a reliable predictor of morphogenesis timing. Altogether, these findings indicate that the timing of key morphogenetic transitions is size-dependent, with larger gastruloids requiring more time to undergo symmetry breaking and axial elongation.

### Size-dependent dynamics of multipolarity in gastruloids

During gastruloid development, axis elongation is coordinated with the differentiation of specialized cell types and the dynamic formation of gene expression patterns. To explore the relationship between morphogenetic events and gene expression, we generated gastruloids from a *Mesp2* reporter line, which express mCherry at the anterior pole [13]. Gastruloids from this reporter line display a similar size-dependent relationship in morphogenesis timing (Fig. S2A-C).

Using high-throughput time-lapse fluorescent imaging, we monitored the spatiotemporal dynamics of *Mesp2*-mCherry expression. We developed a method to identify local peak intensities, allowing us to distinguish between single and multiple *Mesp2* poles (Fig. S2D). Smaller gastruloids (*N*_0_ ≤ 300) consistently exhibit a single *Mesp2* expression pole (Fig. 2A). In contrast, larger gastruloids (*N*_0_ ≥ 600) initially develop up to four poles (Fig. 2B), with the number of poles increasing with gastruloid size (Fig. 2C). Notably, 100% of the largest gastruloids went through a multipolar phase. Nevertheless, by 144 h, over 97% of gastruloids in the *N*_0_ = 150–1200 range resolve their initial multipolarity and achieve uniaxial elongation (Fig. 2C-D, Table 2).

**Figure 2:**
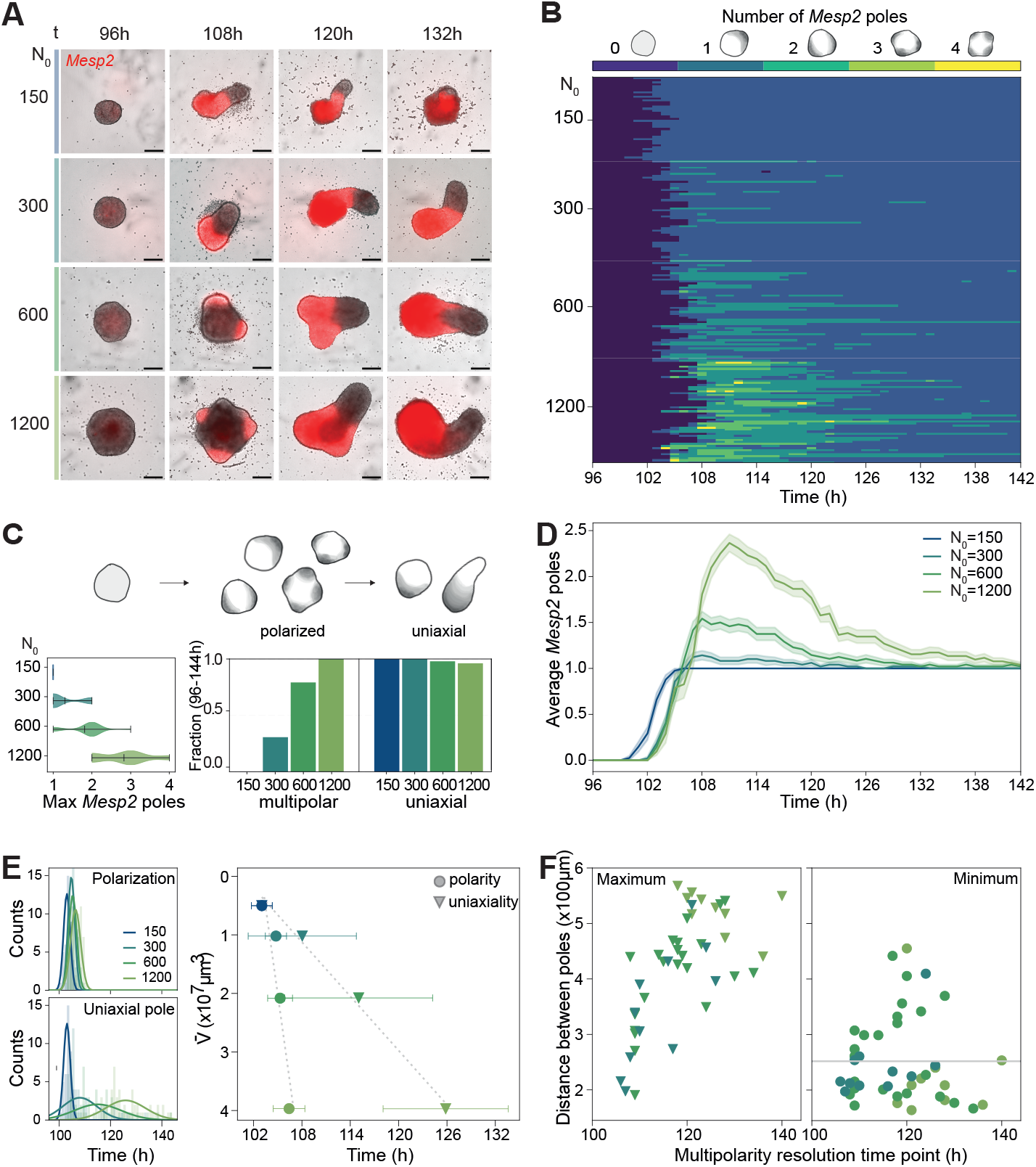
Size-dependent emergence and resolution of multipolarity. (A) Representative images of *Mesp2*-mCherry expressing gastruloids at 72, 96, 120, and 144 h across varying *N*_0_. Scale bars = 200 *µm*. (B) Heatmap of Mesp2-positive pole counts (0 through 4, top legend) over time for gastruloids with *N*_0_ = 150 (n = 41), 300 (n = 49), 600 (n = 48), and 1200 (n = 51). Each vertical line represents a gastruloid (see Supplementary Figure 2D). (C) Cartoon showing unpolarized, polarized, and uniaxial gastruloids; arrows indicate flow of time. Left panel: violin plots of maximum *Mesp2*-positive poles per gastruloid. Right panel: bar plots show the fraction of gastruloids that were multipolar at least once and uniaxial at the end (*>*98%). (D) Temporal dynamics of *Mesp2*-positive poles (mean *±* s.e.m.) for different *N*_0_. (E) Top: histograms of polarization time points (transition from 0 to ≥1 *Mesp2* poles). Bottom: histograms of uniaxial gastruloid formation (transition to one stable *Mesp2* pole). Solid lines indicate Gaussian fits (see Supplementary Tables for detailed statistics). Scatter plot (right) shows mean *±* s.d. of these time points versus gastruloid volume. Gray dotted lines show linear fits to guide the eye. (F) Maximum (left) and minimum (right) distances between peak local maxima of the *Mesp2*-positive poles as functions of multipolarity resolution time (measured as the transition time point from *>*1 *Mesp2* pole to a single stable pole). Solid gray line indicates the average minimum distance across all conditions (min= 253 *±* 77 *µm*).

The timing of polarization, defined as the first detection of an expression pole, is largely consistent across sizes, with only a mean difference of 3 ± 2 h (Fig. 2E, and Table 3). In contrast, the resolution of multipolarity and establishment of a single axis are substantially delayed in larger gastruloids by nearly a day (Fig. 2D-E, Fig. S2D). A strong linear relationship is observed between gastruloid size and the timing of multipolarity resolution, suggesting a decoupling between morphogenetic events, which are size-dependent, and *Mesp2* expression timing, which is robust to size perturbations.

Additionally, the delay in resolving multipolarity in larger gastruloids correlates with increased physical distances between poles (Fig. 2F, Fig. S2B-C). This delay likely reflects the time required to bridge these physical distances during the merging process. Interestingly, the minimum distance between poles is consistent across sizes (Fig. 2F, Fig. S2E).

Previous studies have suggested the existence of an optimal size range for signaling processes mediating symmetry breaking and axial elongation [8, 5, 20]. Our findings support this idea and suggest that exceeding a critical size threshold allows multiple poles to emerge and delays their resolution impeding uniaxial elongation and necessitating longer timescales for axis formation.

Taken together, these results reveal how gastruloid size impacts the timing and robustness of morphogenesis. Surprisingly, despite substantial changes in the timing of key morphogenetic events, the timing of *Mesp2*-mCherry expression remained consistent, suggesting that transcriptional programs are decoupled from morphogenesis dynamics.

### Transcriptional programs are independent of morphogenesis

To test whether size variation affects gene expression, we performed bulk RNA sequencing on gastruloids grown from initial cell numbers of 50, 100, 300, 600, 1200, and 1800 across three experimental batches at 120 h. At this timepoint, gastruloids display striking morphological differences depending on their size (Fig. S3A). As a baseline for significant transcriptional variation, we included a control group of gastruloids (with *N*_0_ = 300) grown without a WNT activation pulse (no-Chiron), previously described failing to elongate or specify germ layers (Van den Brink et al., 2014) (Fig. S3A). Clustering analysis shows a clear separation between treated and no-Chiron samples (Fig. S3B). Principal component analysis (PCA) of the top 1000 most variable genes reveals distinct segregation between no-Chiron controls and gastruloids of varying sizes. These gastruloids are organized along a continuum in PC1 and PC2, which together explain 74% of the variance, suggesting a continuous relationship between size and transcriptional output (Fig. 3A).

**Figure 3:**
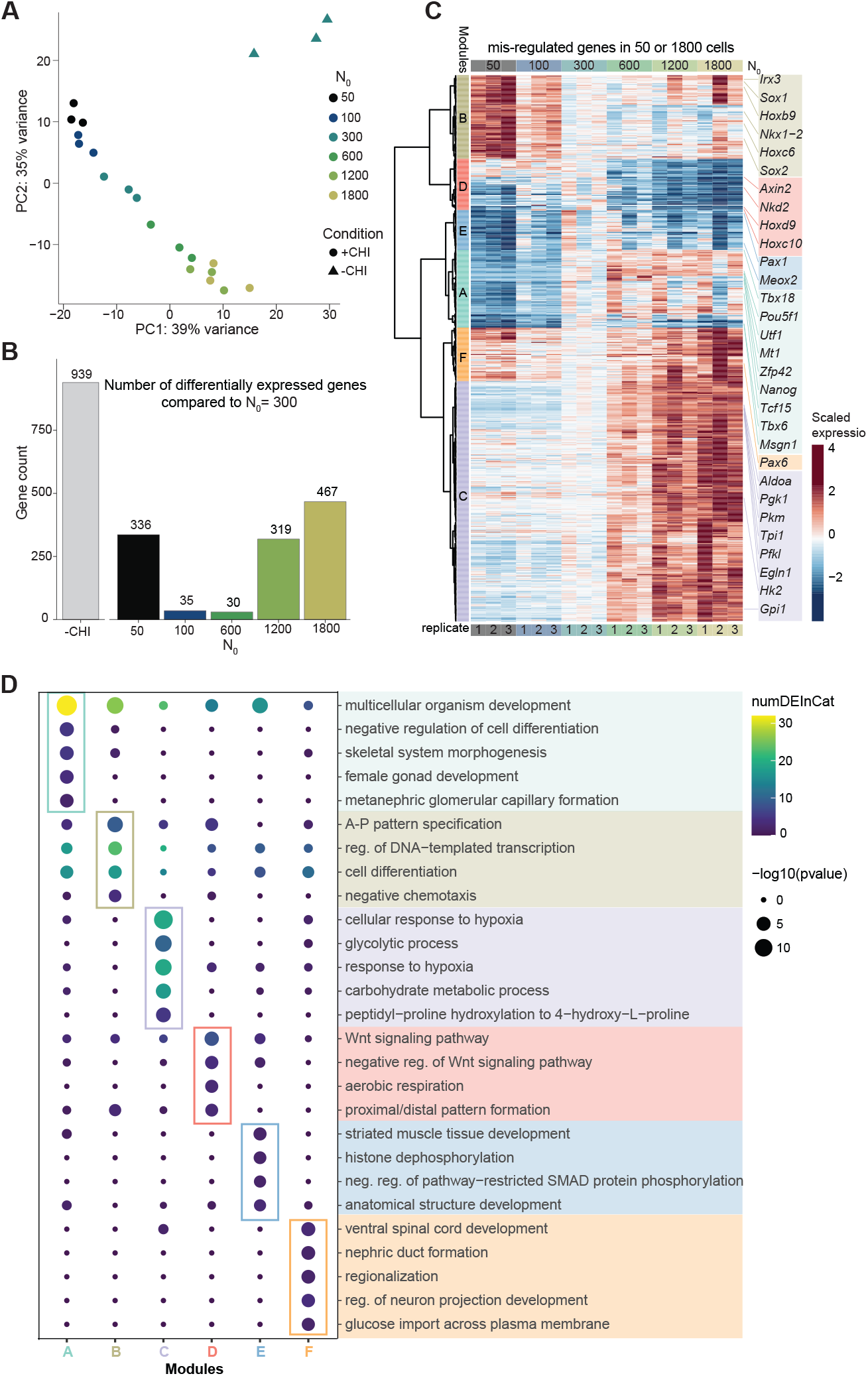
Transcriptional robustness across size variations. (A) Principal component analysis (PCA) of bulk RNA-seq data from gastruloids collected at 120 h. Colors represent *N*_0_ values, and shapes denote treatments (+CHI or no-CHI). Data are from three independent replicates. (B) Bar plot showing the number of differentially expressed genes (p-adj *<* 0.05 and a fold change above 1.5 or below 0.67, measured from DEseq2) for each *N*_0_ relative to *N*_0_ = 300. (C) Heatmap of misregulated genes in extreme sizes (*N*_0_ = 50, 1800), clustered (Ward D2) into six modules (A–F). Expression is scaled across samples. (D) Top five Gene Ontology (GO) terms for each module in (C). Color represents the number of misregulated genes per term; circle size represents the −*log*_10_(p-value) of the GO term enrichment.

Differential expression analysis using *N*_0_ = 300 gastruloids as the reference reveals minimal transcriptional variation across gastruloids seeded from *N*_0_ = 100 to 600, with only 30–35 differentially expressed genes (DEGs) (Fig. 3C). This represents a 30-fold reduction in DEG count compared to the no-Chiron control, highlighting the robust transcriptional output across the *N*_0_ = 100–600 range.

In contrast, extreme sizes (*N*_0_ = 50 and *N*_0_ = 1800) exhibit significant transcriptional changes, though these are still less pronounced than in the no-Chiron control (Fig. 3B, Fig. S3C). We observe progressive transcriptional changes, with smaller gastruloids (*N*_0_ = 100) showing substantial overlap in misregulated genes with *N*_0_ = 50, and upregulated genes in *N*_0_ = 600 largely overlap with those in *N*_0_ = 1200 and 1800 (Fig. S3C).

Focusing on DEGs associated with extreme sizes (*N*_0_ = 50 and *N*_0_ = 1800), we identified six transcriptional modules linked to size. Module A is downregulated and B is upregulated in small gastruloids, whereas module C is upregulated in larger gastruloids, notably in a size-dependent manner (Fig. 3C). Gene ontology analysis shows that modules A and B are enriched for developmental transcription factors, while module C is associated with hypoxia and glycolysis (Fig. 3C-D).

In summary, extreme sizes display morphogenetic changes accompanied by transcriptional shifts. However, within the *N*_0_ = 100-600 range, transcriptional programs remain robust and largely size-independent, despite significant differences in morphogenesis.

### Cell fate composition is robust to size variations

Our bulk transcriptomic analysis reveals that, despite the size-dependence of gastruloid morphogenesis, developmental transcriptional programs are largely conserved across sizes. Specifically, gastruloids within the *N*_0_ = 100–600 range show minimal transcriptional variation, suggesting that cell fate composition remains stable despite striking morphological differences. To investigate this at higher resolution we performed single cell RNA sequencing (scRNAseq) on gastruloids grown from *N*_0_ = 100, 300, 600, 1800 and 5400 cells across two experimental batches at 120 h and 144 h, analyzing a total of 57120 cells (Fig. 4A-B, Fig. S4A-C).

**Figure 4:**
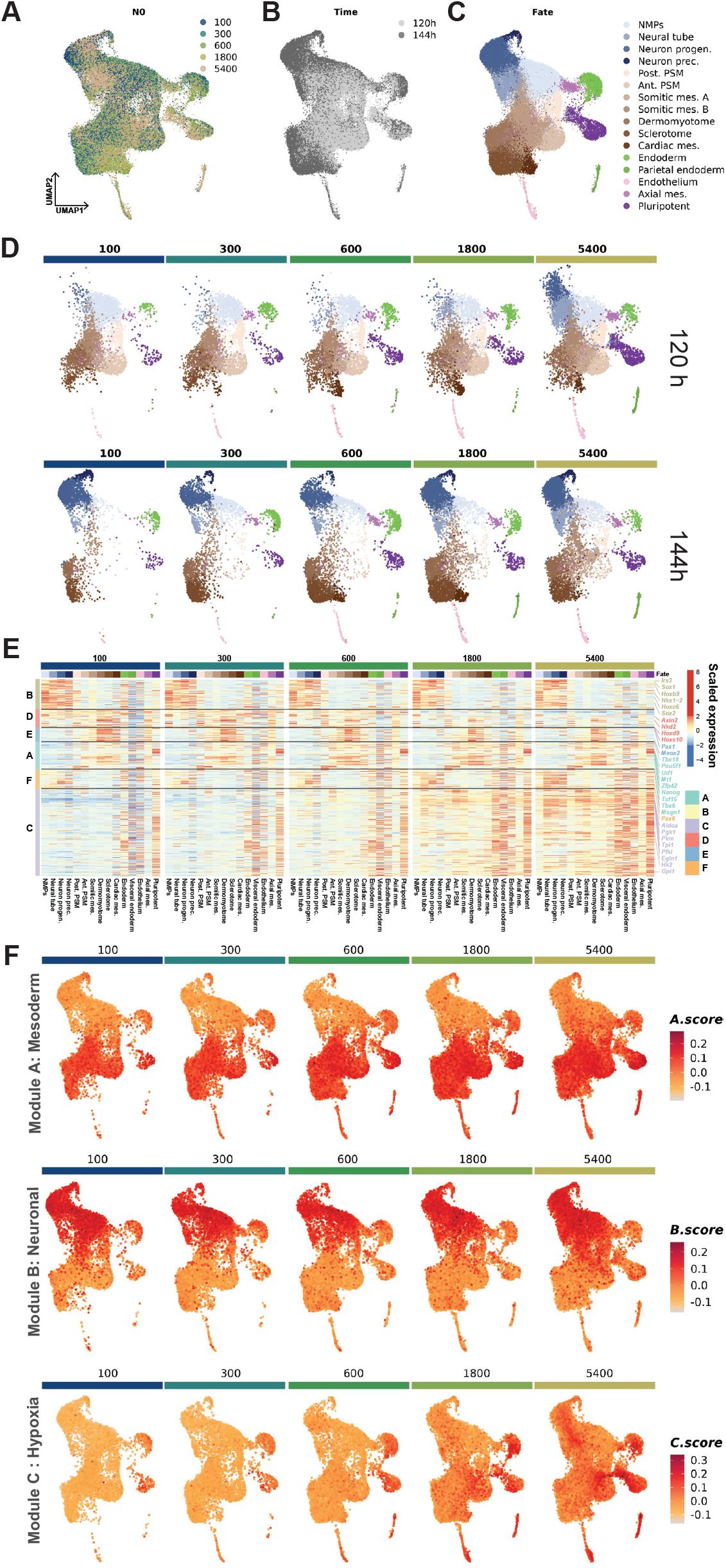
Cell fate composition is robust to size variations. (A-C) UMAP projections of single-cell RNA-seq data (57,000 cells) from gastruloids collected at 120 h and 144 h, colored by *N*_0_ (A), collection time (B), and cell fate (C). (D) UMAP projections split by *N*_0_ and collection time, with color indicating cell fates. (E) Heatmap of pseudo-bulk gene expression for genes misregulated in extreme sizes (*N*_0_ = 50 and 1800) from Fig. 3C. Expression is scaled across the entire dataset, with separate panels for each *N*_0_. (F) UMAP projections of single-cell RNA-seq data split by *N*_0_ and collection time, with cells colored by scores for modules A, B, and C (from Fig. 3C).

Expected lineages corresponding to the three germ layers (mesoderm, ectoderm, and endoderm) are observed (Fig. 4C), consistent with previous studies [18, 17, 13]. At 120 h, gastruloids from *N*_0_ = 100–1800 cells exhibit similar cell compositions, with batch-to-batch variation exceeding size-dependent variation (Fig. 4D, Fig. S4D-E). However, smaller gastruloids display a higher proportion of neuro-mesodermal progenitor cells (NMPs). NMPs have the potential to differentiate into both neuronal or mesodermal lineages and are essential for embryonic axial elongation [23, 24, 17, 18, 13]. At 144 h, larger gastruloids (from *N*_0_ = 300–1800) remain highly similar (batch-to-batch differences higher again), while *N*_0_ = 100 gastruloids show depleted progenitor pools, particularly NMPs and pre-somitic mesoderm (PSM). This depletion is particularly prominent in batch 1, where all *N*_0_ = 100 gastruloids collapsed by 144 hours (Fig. S4B), suggesting that the collapse is linked to progenitor cell exhaustion. Lineage marker gene expression remains consistent across size variations (Fig. S4F), indicating that transcriptional programs are properly established even under severe morphological alterations. To dissect whether transcriptional changes observed in bulk RNAseq for extreme sizes (*N*_0_= 50, 1800) are due to shifts in cell populations or intrinsic gene expression, we generated pseudobulk measurements for each cell fate across sizes. Most transcriptional modules (A,B,D,E,F) exhibit limited size-dependent variation and instead reflect proportional changes in cell proportion (Fig. 4E). Module A, downregulated in small gastruloids (Fig. 3D), is associated with mesodermal and pluripotent lineages. Module B, upregulated in smaller gastruloids, corresponds to neuronal lineages (Fig. 4F). In contrast, module C, associated with hypoxia and glycolysis, showed a unique size-dependent response, being upregulated across all lineages in larger gastruloids, particularly in endodermal, endothelial, and pluripotent lineages (Fig. 4E-F).

Overall, aside from a neuronal-to-mesodermal bias in smaller gastruloids and a size-dependent increase in glycolysis and hypoxia responses, our findings indicate that gastruloid transcriptional status and cell fate composition are largely decoupled from size and morphology.

### Effective system size governs morphogenesis and patterning

Our results demonstrate that size variations affect the timing and outcomes of morphogenesis, while transcriptional states and cell fate composition remain largely conserved. However, at extreme sizes (*N*_0_ ≤ 100 or *N*_0_ ≥ 1200), morphogenetic changes are accompanied by transcriptional shifts. To explore how system size influences these processes at size boundaries, we conducted size perturbation experiments to assess whether gastruloid behaviors could be rescued by altering physical dimensions mid-development.

We hypothesized that gastruloids might retain memory of their initial seeding number, with size-dependent system properties established early in development influencing subsequent morphogenesis. Indeed, our scRNA-seq data reveal differential hypoxic responses correlating with gastruloids size, suggesting that metabolic processes sensitive to initial cell number may propagate through development in a size-dependent manner. Alternatively, morphogenesis could be a purely size-dependent process, with gastruloid behavior determined by the effective cell number at a given time point, irrespective of initial seeding conditions.

To test these hypotheses, we developed a method to manipulate gastruloid size at 72 h, immediately following the WNT activation pulse when gastruloids are spherical. Microsurgical manipulations included fusing smaller gastruloids or cutting larger gastruloids to match reference sizes, e.g. *N*_0_ = 300 (Fig. 5A). Post-manipulation measurements confirm the validity of this approach: fused *N*_0_ = 50 gastruloids (50×6) and cut *N*_0_ = 1200 gastruloids (1200/4) match the size of *N*_0_ = 300 controls (Fig. 5B-D, Fig. S5A-C).

**Figure 5:**
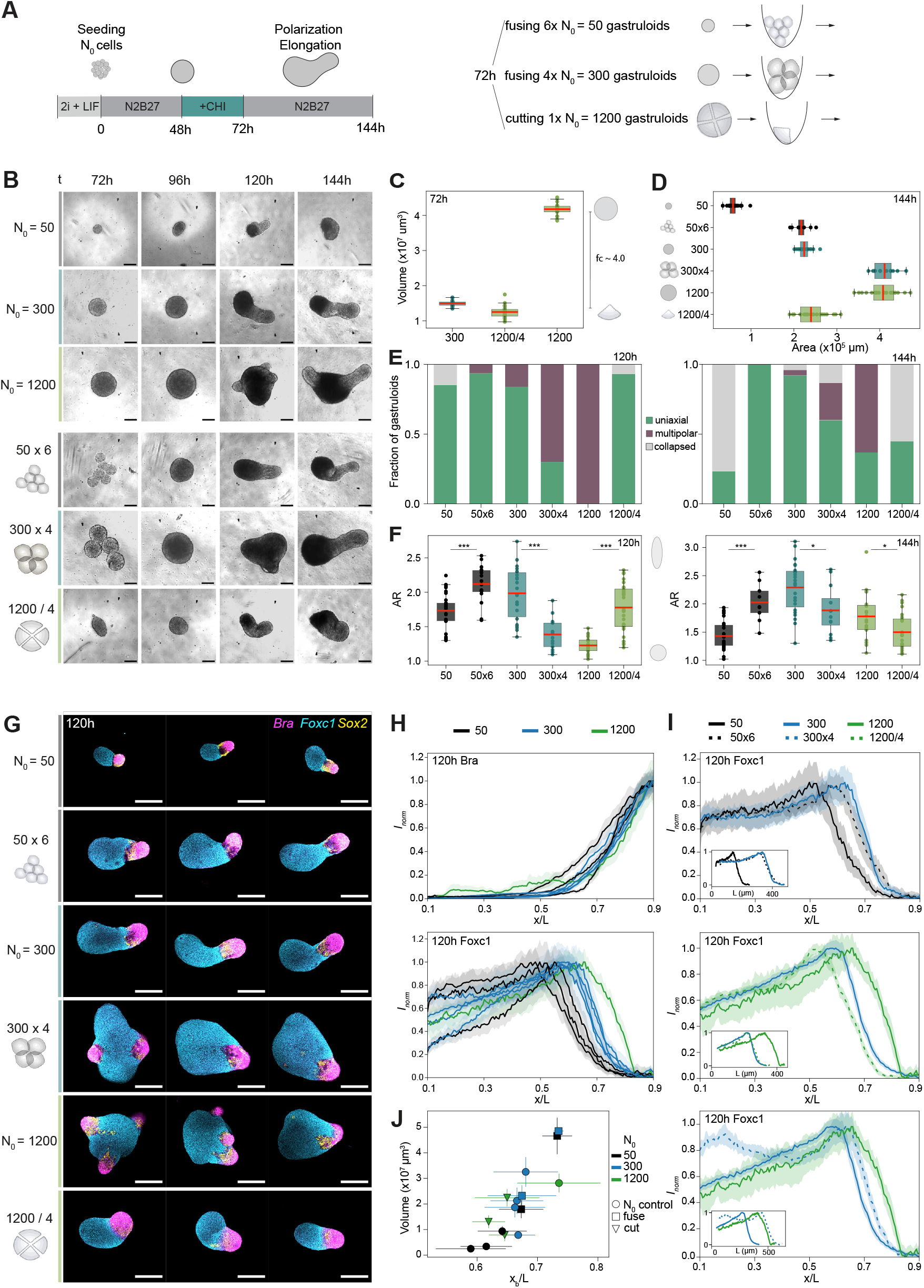
Resizing gastruloids rescues morphogenesis and patterning. (A) Gastruloid size perturbation protocol. At 72 h, gastruloids were manipulated by fusion (6x *N*_0_ = 50 and 4x *N*_0_ = 300, “fused”) or dissection (*N*_0_ = 1200 into 4 pieces, “cut”). Perturbed gastruloids were grown under standard protocol conditions until 144 h. (B) Brightfield images of gastruloids at 72, 96, 120, and 144 h post-seeding, showing morphology across varying *N*_0_ for control, fused, or cut conditions. Scale bars = 200 *µ*m. (C) Mean gastruloid area after perturbation at 72 h for *N*_0_ = 300, *N*_0_ = 1200, and cut *N*_0_ = 1200 (1200/4). Boxplots show group mean, with whiskers extending to the farthest data point within 1.5x the interquartile range (IQR). Fold change (fc) for *N*_0_ = 1200 : 1200/4 is indicated. (D) Mean gastruloid area at 144 h across control, cut, and fused gastruloids. Boxplot characteristics as in C. (E) Proportions of gastruloids categorized as uniaxial, multipolar, or collapsed for control and perturbed conditions at 120 h (left) and 144 h (right), based on manual annotation. (F) Gastruloid aspect ratio at 120 h (left) and 144 h (right), used as a proxy for uniaxial elongation. Boxplot characteristics as in D. P-values for control vs. perturbed conditions were calculated using a two-sided independent t-test: p *<* 0.05 (*), p *<* 0.01 (**), and p *<* 0.001 (***); n.s. = not significant. (G) Maximum projections of confocal image stacks of 120 h gastruloids immunofluorescently stained for Bra, Foxc1, and Sox2. Posterior ends are oriented to the right. Scale bars = 200 *µ*m. (H) Normalized expression profiles (means *±* s.e.m.) of Bra (top) and Foxc1 (bottom) at 120 h for gastruloids with the same *N*_0_, plotted as a function of the relative position (x/L) along the average midline. Data from four experimental batches are shown. AP-axis is oriented left-to-right. (I) Expression profiles of Foxc1 at 120 h, comparing perturbed (dashed lines) and control (solid lines) conditions. Insets show normalized mean expression profiles as a function of average absolute position. (J) Scatter plot of mean pattern boundary positions (xb, half-maximal expression) of Foxc1 versus mean gastruloid volumes for each condition across four batches. Error bars represent s.d. of xb and volumes.

Smaller gastruloids (*N*_0_ = 50 cells) typically elongate by 96 h and collapse by 144 h (Fig. 5B-E, Fig. S5D). When fused to achieve *N*_0_ = 300, however, these gastruloids no longer exhibit collapse, suggesting that collapse results from progenitor cell exhaustion and depends on effective size. Similarly, cutting *N*_0_ = 1200 gastruloids into smaller fragments approximating the *N*_0_ = 300 reference size rescues the multipolarity phenotype, with most gastruloids achieving uniaxial elongation by 120 h (Fig. 5B-E-F, Fig. S5D-E). Nonetheless, cutting introduces variability in fragment sizes, with smaller fragments prone to collapse (Fig. 5E, Fig. S5D). Conversely, fusing *N*_0_ = 300 gastruloids into a larger gastruloids increases multipolarity and reduces elongation (Fig. 5B-F, Fig. S5D-E). These findings suggest that gastruloid morphogenesis is primarily governed by the effective size rather than initial seeding number.

Although transcription drives cell fate decisions, morphogenetic movements organize cells into functional domains, contributing to the spatial order observed in gastruloids. To examine how size influences transcriptional states (at the cellular level) and global morphology (at the tissue level), we analyzed gene expression patterns using immunofluorescence staining for germ-layer markers Bra, FoxC1 and Sox2 (Fig. 5G, Fig. S5I). Maximum intensity projections of confocal stacks were used to extract 1D intensity profiles along each gastruloid’s midline (see Methods).

In line with earlier observations, smaller gastruloids (*N*_0_ ≤ 300) consistently exhibited a single posterior Bra-positive pole, while larger gastruloids displayed multiple Bra-positive poles (Fig. 5G, Fig. S5I). Across four experimental batches, normalized Bra profiles show greater batch-to-batch variation than size-dependent differences (Fig. 5H, S5F-G). In contrast, FoxC1 exhibits a size-dependent trend, with larger gastruloids showing an expanded FoxC1-positive domain at 120 h. By 144 h, FoxC1 expression patterns became more uniform across sizes, though with increased variability (Fig. S5I-J). These results suggest that size thresholds for pattern formation may differ among distinct cell and tissue types: Bra patterns are consistent across sizes, whereas FoxC1 patterns are size-dependent.

To determine whether gastruloids adapt gene expression patterns to new sizes after perturbation, we compare FoxC1 expression between microsurgically manipulated and *N*_0_ sized gastruloids. Remarkably, FoxC1 patterns aligned with the new sizes rather than the original seeding size (Fig. 5I). There was no correlation between initial seeding size and pattern boundary positions (xb) (Fig. S5I). Instead, normalized pattern boundary positions xb/L strongly correlated with effective gastruloid size (Fig. 5J, Fig. S5H). These findings reveal that gastruloids adapt both morphologically and in terms of patterning to their new size, demonstrating developmental plasticity.

Overall, these findings highlight that gastruloids morphogenesis and patterning are governed by physical properties such system size rather than fixed developmental programs. As gastru loids reach specific size thresholds, emergent features such as multipolarity and expanded expression domains deviate from simple scaling behaviors. This underscores the modularity of self-organizing systems, revealing that gastruloids produce predictable outcomes that depend on effective size.

## Discussion

Our findings reveal a surprising temporal decoupling of transcriptional programs from morphogenetic events, offering new insights into developmental complexity. This decoupling is size-dependent, with physical parameters such as system size and cell number governing morphogenetic dynamics. While reaction-diffusion mechanisms have been proposed to explain symmetry breaking, previous studies highlight the critical role of cell adhesion in organizing gastruloid morphogenesis and patterning. For instance, differential adhesion was proposed to drive endoderm organization [25] and arrange Wnt activity domains into a single pole defining the anteroposterior axis [14]. Gastruloid elongation similarly aligns with convergent extension, driven by active cell crawling and differential adhesion [26, 13].

Our observations suggest that adjusting system size may influence cell sorting dynamics and tissue rearrangements. Larger gastruloids exhibit delayed symmetry breaking, increased multipolarity, and prolonged elongation, suggesting that system size controls the timing and merging of morphological poles. Conversely, smaller gastruloids display accelerated morphogenesis but often collapse, linking size constraints to progenitor pool exhaustion.

Despite the striking morphological phenotypes observed across a broad size range, cell fate composition remains stable. However, extreme sizes additionally trigger metabolic shifts. Hypoxia and glucose metabolism are known to regulate differentiation in gastruloids and embryos [27, 28, 29]. Hypoxia enhances spontaneous elongation and lineage representation [30], while glucose metabolism biases differentiation toward neuronal or mesodermal lineages [19].

In larger gastruloids, pluripotent cells localize to an inner core [20], restricting oxygen availability and intensifying hypoxic responses. Conversely, the absence of hypoxia in smaller gastruloids accelerates differentiation, rapidly depleting progenitor pools. Additionally, the smaller absolute number of progenitors in small gastruloids may explain their collapse, as progenitor states are depleted earlier due to simple numerical constraints.

Our resizing experiments demonstrate that morphogenesis does not depend solely on transcriptional states or initial cell fates but instead arises from emergent physical properties. By manipulating gastruloid size mid-development, we reveal that morphogenetic trajectories adapt to effective size rather than retaining memory of initial seeding conditions.

This ability to “reset” developmental processes highlights a fundamental plasticity in multicellular systems. Physical parameters such as size and cell number, rather than early developmental memory, govern morphogenetic transitions. This finding underscores the emergent nature of tissue organization and reveals the gastruloid system’s capacity to reorganize morphogenetic outcomes dynamically.

Remarkably, transcriptional programs and cell fate composition remain robust across a wide size range, even when morphogenesis is significantly altered. This stability highlights a decoupling between physical constraints and gene regulatory networks, ensuring consistent cell fate decisions despite size-induced variability. At extreme sizes, however, distinct transcriptional modules emerge, associated with hypoxia and glycolysis. The size-dependent activation of metabolic pathways suggests that hypoxia and glycolysis act as integrators of physical constraints and developmental regulation. This adaptability may represent a mechanism to buffer size perturbations while maintaining overall developmental trajectories.

The temporal decoupling of transcriptional programs from morphogenetic events may facilitate evolutionary change. This extra degree of freedom in morphogenesis from gene regulatory networks could enable the emergence of new forms and structures without disrupting core developmental programs.

Unlike embryos, which have extra-embryonic tissues and where size is tightly regulated [31, 32, 33], gastruloids lack these constraints. As a result, they can explore a broader range of morphological states, generating variability that is adaptively accommodated. This plasticity underscores the modularity of self-organizing systems, where robust developmental outcomes emerge from simple physical parameters such as size and cell number.

Our findings highlight the versatility of gastruloids as an experimental model. The robustness of transcriptional states and cell fate composition to size variations makes gastruloids amenable to diverse experimental approaches. Smaller gastruloids are ideal for high-resolution microscopy, while larger ones provide sufficient biological material for biochemical and molecular assays. This flexibility positions gastruloids as a powerful platform for studying spatiotemporal dynamics in mammalian development across varying experimental scales. By providing a simplified, self-organizing system, gastruloids allow us to disentangle physical, mechanical, and biochemical contributions to developmental processes.

The observed temporal decoupling of transcriptional states and morphogenetic events opens new avenues to investigate the integration of biochemical cues, mechanical forces, and geometric constraints across spatial and temporal scales. Understanding these mechanisms will advance our knowledge of how robust morphologies arise during development and may inform strategies for engineering self-organizing tissues in regenerative medicine and bioengineering. By uncovering the fundamental principles of developmental plasticity and physical constraints, gastruloids offer unique opportunities to address central questions in developmental biology and tissue engineering.

## Methods

### mESC culture

129/SvEv (EmbryoMax) mouse embryonic stem (mES) cells were cultured on gelatin-coated six-well plates in a humidified incubator (5% CO_2_, 37°C). Cells were maintained in LIF + 2i DMEM medium composed of: DMEM 1X + Glutamax (Fisher 11584516) supplemented with 10% Decomplemented FBS (Gibco, 11573397, decomplemented 30 min at 56°C), 1X Non-essential amino acids (NEAA, Gibco 11140-035), 1mM Sodium Pyruvate (Gibco,11360-039), 1% Penicillin-Streptomycin (Gibco, 15140-122), 100 *µ*M 2-Mercaptoethanol (Gibco 31350-010),10 ng/mL Leukemia Inhibitory Factor (LIF, Miltenyi Biotec 130-099-895), 3 *µ*M GSK3 inhibitor CHIR 99021 (Chiron, Sigma, SML1046), 1 *µ*M MEK inhibitor PDO35901 (Sigma, PZ0162). Experiments were performed using cells between passages 20 and 30. Cells were passaged every other day as follows: cells were washed with PBS (Gibco, 10010023) and dissociated using Trypsin (Sigma T3924) or Accutase© (StemPro Ref: A11105-01). Detached cells were resuspended in DMEM, counted using an automatic cell counter (Logos Biosystems LUNA-II) and reseeded at a density of 200,000–400,000 cells per well.. When cells were not passaged, half of the culture medium was replaced. Cells were tested regularly for mycoplasma contamination using the Eurofins MycoplasmaCheck service.

### Gastruloid culture

Gastruloids were generated as previously described in [10]. N2B27 medium was prepared in-house every three weeks using the following components: 250 mL DMEM/F12+GlutaMax (Gibco, 10565018), 250 mL Neurobasal (Gibco, 21103049), 2.5 mL N2 (Gibco, 17502-048), 5 mL B27 (Gibco, 17504-044), 1X Non-Essential Amino Acids (NEAA, Gibco, 11140-035), 1 mM Sodium Pyruvate (Gibco, 11360-039), 100 *µ*M 2-Mercaptoethanol (Gibco, 31350-010), 1% Penicillin-Streptomycin (Gibco, 15140-122), and 2.5 mL GlutaMax (Gibco, 35050061). Initial cell seeding was performed manually using a multipipette and the cell counts were determined with an automatic cell counter (Logos Biosystems LUNA-II). Gastruloid experiments were performed in three laboratories using two slightly different protocols. For cell dissociation, cells treated with Accutase were immediately resuspended in N2B27 medium. When Trypsin was used for dissociation, cells were rinsed twice with phosphate-buffered saline (PBS) before resuspension in N2B27. The dissociated cells were seeded into Costar Low Binding 96-well plates (Costar, Corning, 7007) at a volume of 40 *µ*L per well. After 48 hours of aggregation, the spheroids were subjected to a 24-hour pulse of Wnt agonist by adding 150 *µ*L of 3 *µ*M CHIR 99021 (Chiron) in N2B27 to each well, unless otherwise specified. Subsequently, 150 *µ*L of the medium was replaced every 24 hours until gastruloid collection.

### Generation of mutant ES cells by CRISPR/Cas9

Wild-type mESCs (EmbryoMax 129/SVEV) were used to generate a cell line heterozygote for the *Mesp* locus using the CRISPR/Cas9 genome editing protocol described in [34]. Then, we integrated reporter constructs consisting of p2a-EGFP-NLS-PEST and p2a-mCherry-NLS-PEST in frame with the *Mesp1* and *Mesp2* coding sequence, respectively. We used a template repair knock-in strategy using a mini pUc57 plasmid containing the reporter constructs surrounded by homology arms targeting either the *Mesp1* or the *Mesp2* coding sequence. These template repair plasmids were co-transfected with a single guide RNA (sgRNA) Cas9 plasmid. ES cells were transfected with 5 *µ*g of sgRNA-Cas9 plasmid (and 1.5 *µ*g of “reporter plasmid” when applicable) using the Promega FuGENE 6 transfection kit and dissociated 48h later for puromycin selection (1.5 *µ*g/ml). Clone picking was done 5–6 days later, and positive ES cell clones were assessed by PCR screen using the MyTaq PCR mix kit (Meridian Bioscience) and specific primers surrounding the targeted region (Table 4). Mutations were verified by Sanger sequencing. The region to be deleted were targeted by two flanking sgRNA for deletions, and one sgRNA for the integration of reporter constructs. all guides are listed in Table 5) sgRNAs were designed using the CRISPR Guide RNA Design Tool from Benchling. sgRNA sequences were inserted in a Cas9T2APuromycin expressing plasmid containing the U6 gRNA scaffold (“sgRNACas9 plasmid”, gift of A. Németh; Addgene plasmid, 101039).

### Size perturbation: cutting and fusing gastruloids

Gastruloid size perturbation was performed at 72 h post seeding, right after the Chiron pulse. To fuse multiple smaller gastruloids (6x *N*_0_ = 50 and 4x *N*_0_ = 300), gastruloids were collected and pooled in a 60 mm Petri dish with pre-warmed N2B27 medium. The respective number of gastruloids were collected using a cut and coated P200 pipette tip and transferred into a well of a new ultra-low binding 96well U-bottom dish. For resizing larger gastruloids to multiple smaller ones (*N*_0_ = 1200/4), a gastruloid was transferred to a Petri dish with pre-warmed N2B27 medium and first cut in half and then each half was cut into a quarter using a tungsten needle. Each quarter gastruloid was subsequently transferred into a separate well of a new ultra-low binding 96well U-bottom dish as described above. Gastruloid dissection was designed to minimize tissue loss and maintain equal proportions of each tissue part, however total cell recovery and optimal tissue quarters were imperfect, as reflected in the volume measurements comparing the conditions *N*_0_ = 300, *N*_0_ = 1200, and *N*_0_ = 1200/4 condition (Fig. S5B). Fusion and cutting procedures each took a few minutes up to an hour, depending on the sample number.

### Immunofluorescent staining

Gastruloids at 120 h and 144 h were collected from the well plates, pooled in a 15 ml falcon tube, and washed once with PBS with Mg2+ and Ca2+ (PBS++, Gibco, 14040133). Gastruloids were subsequently fixed in 10 ml 4% paraformaldehyde solution (PFA, Thermo Scientific Chemicals, 30525-89-4) for 2 h, afterwards washed twice with 10 ml PBSF (10% FBS in PBS++), resuspended in 1 ml in PBS++ and stored at 4°C (for several weeks). For the immunofluorescent staining, gastruloids were first permeabilized in 10 ml PBSFT (10% FBS and 0.03% Triton in PBS++) and incubated for 1 h at room temperature (RT). Gastruloids were then incubated in 0.5 ml PBSFT containing 4’,6-diamidino-2-phenylindole (DAPI) and primary antibody over night at 4°C (see Table 6 for details on antibodies and concentrations). On the next day, gastruloids were washed three times with 10 ml PBSFT at RT for 30 min each and subsequently incubated in 0.5ml PBSFT containing DAPI and secondary antibody over night at 4°C (Table 6). Gastruloids were washed twice in 10 ml PBSFT and once in PBS++ at RT for 30 min each. All washes and incubations were performed under nutation. For the mounting procedure, all access PBS++ was removed from the tube and replaced by 200 *µ*l mounting medium composed of 50:50 Aqua-Poly Mount (Polysciences 18606-20) and PBS++. Gastruloids in mounting medium were then transferred to a round glass bottom dish, covered with a cover glass and nail polish sealed.

### Brightfield imaging OlympusCKX41

To record the morphological development of gastruloids following size perturbation, brightfield (BF) images of each gastruloid in the U-bottom well were taken using an Olympus CKX41 inverted phase-contrast microscope. Images were collected every 24 h from 72-144 h post seeding at a 10x magnification.

### Confocal imaging

Confocal fluorescent imaging of fixed and stained gastruloids was performed on a Zeiss LSM880 and a Zeiss LSM980 confocal microscope. Gastruloids were imaged individually using a Zeiss 10X, 0.3 numerical aperture air objective, and a 150 *µ*m-thick z-stack of 30 slices with a voxel size of 1.186 × 1.186 × 5.000 *µ*m^3^. Laser lines 405 nm, 488 nm, 561 nm, and 633 nm were used to image DAPI, AF-488, AF-546, and AF-647, respectively. Confocal images of gastruloids were used to extract morphological parameters and 1D gene expression profiles.

### Live movie image analysis

Live movies were acquired using an Olympus video-microscope with the Olympus CellSens dimension 3.1 software, equipped with a Hamamatsu C11440-36U CCD camera with a pixel size of 5.86 × 5.86 *µ*m and a 4X 0.13 NA objective or an IncuCyte S3 (Sartorius) microscope with a 10X objective and 400 ms exposition for the red channel. BF and fluorescent images of individual gastruloids in the 96 U-bottom well plates were taken every hour for several days of gastruloid development.

### Morphological analysis

Gastruloid segmentation was performed on BF images using a SegmentAnything Model from Meta AI [35]. Gastruloid masks were then used to calculate the gastruloid contour from which the perimeter *P* and area *A* were derived. The aspect ratio (*AR*) of a gastruloid was determined by fitting an ellipse to the extracted whole gastruloid mask (`skimage.measure.regionprops` function) and taking the ratio of the major-to-minor axis length, therefor increasing with gastruloid AP axis elongation. The circularity of a gastruloid was calculated from the extracted perimeter *P* and 2D projected area *A*

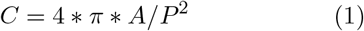

and is thus defined as a measure between 1 (perfect circle) and 0, which decreases as gastruloids lose their spherical morphology. For time points where the gastruloid morphology was approximately spherical, the gastruloid volumes *V* were reconstructed using the area measurement *A*

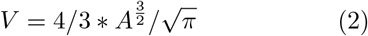

### Optimal partitioning

To determine time points of morphological transition within time series data, optimal partitioning was used to classify the data into two statistically distinct segments *s*_1_ and *s*_2_ by minimizing the cost function *C*

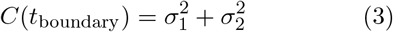

where 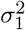 and 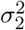 are the inter-segment variances of *s*_1_ and *s*_2_, respectively, if the data is partitioned at time point *t*_boundary_. Sweeping through all feasible *t*_boundary_ values lets us identify the time point that minimizes *C*

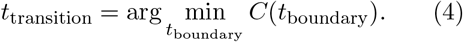

which defines the point in time where the morphological measurements transition from one relatively homogeneous regime to another and therefore effectively capturing a change in the underlying dynamic morphogenetic processes (e.g., symmetry breaking or morphological elongation).

### Mesp2 pole quantification

The quantification of anterior pole dynamics was performed on fluorescent one-hour interval live movies of Mesp2 gastruloids. First, each fluorescent image was filtered using a median rank filter with kernel size k=7 to reduce salt and pepper noise. The filtered fluorescent image was then masked using the gastruloid contour extracted from the BF image and a binary mask was computed using a threshold optimized across all size conditions, time points and to reduce autofluorescence detection. Because the Mesp2 fluorescent signal is often non-uniform within a single anterior pole, especially for time points *>*120 h, an additional smoothening of the fluorescent image was performed using a Gaussian filter (sigma=20) to avoid counting multiple poles resulting from this artifact. The smoothed image was subsequently masked with the binarized fluorescent image and the peak local maxima were determined. Each maximum defines an anterior Mesp2 pole. For every gastruloid, it was thus possible to extract a count of anterior Mesp2 poles at each imaged time point.

### Distance between Mesp2 poles

Mesp2 pole distances were obtained by calculating the Euclidian distance *D* between the point coordinates (*x, y*) of any two peak local maxima detected in the fluorescent image

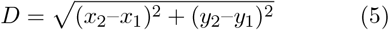

### Confocal image analysis

For each confocal imaging stack, i.e. a single gastruloid, 2D maximum projections were computed of all channels and used to extract morphological parameters and 1D gene expression profiles largely following the analysis outlined in [11] (link to github).

### Morphological analysis

In brief, the DAPI channel was used to mask and determine the gastruloid contour, and to define the major body axis, by computing the medial axis and extrapolating at each end to a point on the gastruloid contour. The intersection between the medial axis ends and the gastruloid contour define the anterior and posterior tips of a gastruloid and the total length of the extrapolated medial axis defines gastruloid length L. Reconstruction of gastruloid volumes was achieved by calculating *n*_*b*_=200 non-overlapping segments of the extrapolated medial axis, equidistantly spaced along each side of the gastruloid contour. Assuming radial symmetry of a gastruloid along its major body axis, the volume of each segment can be approximated and the sum across all segments defines gastruloid volume *V*.

### 1D Gene expression profile analysis

Only gastruloids for which an unbranched medial axis could be defined were considered for gene expression profile analysis. Morphological segmentation along the medial axis into *n*_*b*_ bins, was used to compute an average maximum projection fluorescent intensity *I* over each bin, for every channel. To facilitate profile comparison across different experimental batches and conditions, each gastruloid intensity profile was normalized by their condition mean

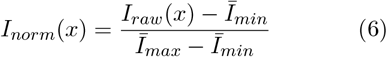

Normalized intensity profiles were plotted as a function of the position along the gastruloid’s major body axis with length *L* for 0.1 *L* 0.9 or of the fractional position *x/L* for 0.1 *x/L* 0.9. Gene expression profile boundary positions *x*_*b*_*/L* are defined as the fractional positions along the gastruloid midline where the half-maximal expression level within the boundary regions is reached.

### Bulk-RNA barcoding and sequencing (BRB-seq)

Gastruloids seeded from different cell numbers (50, 100, 300, 600, 1200, 1800 cells with the regular Chiron treatment and 300 cells without the addition of Chiron) were grown until 120h as described above. For each replicate (three in total), 60 gastruloids from each condition were collected in a 1.7ml Eppendorf tube and washed once with PBS, pelleted and kept in −80°C until all samples were collected. Each sample were then thawed and extracted using the RNeasy mini kit (Qiagen) according to manufacturer’s recommendation with on-column DNase digestion. RNA were quantified using Qubit Fluorometric Quantification and RNA quality was assessed using a TapeStation TS4200. All samples had an RNA integrity number (RIN) above 9. Libraries were performed using Alithea Genomics Mercurius protocol v.0.2.2 and sequenced on Novaseq 6000 in a PE run with 28, 8i, 90 configuration.

### Bulk-RNA sequencing (BRB-seq analysis)

Reads were assigned to genes and to samples using STAR Solo [36] version 2.7.10b `(--soloStrand Forward –soloType CB UMI Simple --soloCellFilter None --soloFeatures Gene)`. The GTF file used for gene annotation is available on Zenodo [37].

All counts were aggregated into a single matrix, and a second matrix was generated where the counts were normalized to the million reads. The normalized counts were used to compute PCA and clustering. Only the 1000 genes with the highest variance were kept. Only protein-coding genes were considered for the differential gene expression analysis computed with DESeq2 [38] using the Wald test. A gene was considered differentially expressed if the adjusted *p*-value was below 0.05 and the fold-change was above 1.5 (or below 0.67).

The Euler diagrams were generated with the `eulerr R` package [39]. The modules were identified using genes differentially expressed between the control condition and 50 cells or between the control condition and 1800 cells. The normalized expression of each of these genes across all samples with Chiron was scaled to achieve a standard deviation of 1 and a mean of 0 in the samples with 300 cells.

The genes in this scaled matrix were clustered using Pearson’s correlation between genes and the ward.D2 algorithm. Six groups were obtained by cutting the clustering tree. Gene Ontology analysis was performed using the genes of each module with the goseq R package [40].

### Single cell-RNA-sequencing (scRNA-seq)

scRNA-seq was performed as previously described [13]. Briefly, gastruloids seeded from different cell numbers (100, 300, 600, 1800, 5400 cells) were grown until 120 h and 144 h across two independent replicates. For each condition the number of gastruloids used was chosen to unsure that a minimum of 200 000 cells were obtained for each sample and a minimum of 24 gastruloids were collected for each sample to limit the impact of gastruloids-to-gastruloids variation. Gastruloids were collected and washed in 1ml of PBS in a 1.7 ml Eppendorf tube and dissociated using 100*µ*l of Accutase (Stempro) for 5 minutes at 37°C. Full dissociation was verified to ensure absence of doublets and if necessary, it was completed using mechanical dissociation by pipetting. All centrifugation were done at 400 g for 5 minutes. Conditions were multiplex using the CellPlex procedure according to manufacturer’s recommendations. Cells were incubated in 50*µ*l of cell multiplexing oligos (3’ CellPlex Kit Set A, PN-1000261) for 5 minutes at room temperature. They were then thoroughly washed three times with 1 ml PBS 1% BSA ensuring to remove as much as possible of the supernatant each time to prevent sample-to-sample contamination. Each sample was then counted and viability was assessed using a Countess 3 automated cell counter (Invitrogen) and viability was above 90% in all cases. Samples were then pooled in desired proportion to ensure proper representation of each experimental condition and the pooled cell suspension was filtered using a 40 *µ*m cell strainer (Flowmi, BAH136800040). The final count was performed and 24 000 cells were targeted for recovery using the 10x Genomics approach following their recommendations since multiplexing allows for the resolution of more doublet cells, yielding on average 15000 singlet cells that can be used for analysis. cDNA preparations were performed according to 10x Genomics recommendations and were amplified for 10-12 cycles and cDNA libraries were assessed on fragment analyser. Both cell multiplexing oligo and gene expression libraries were generated according to 10x Genomics reccomendations and were sequenced on a Novaseq (Illumina protocol #1000000106351 v03) with the cbot2 chemistry.

### scRNA-seq analysis

Single-cell analysis was performed as previously described [13]. Fastq files containing the sample information (cell multiplexing oligo) were processed with CITE-seq-Count version 1.4.4 [41] using the following arguments:

~~~
--cell_barcode_first_base 1
--cell_barcode_last_base 16
--umi_first_base 17 --umi_last_base 28
--expected_cells 24000 --whitelist
‘cellranger_barcodes_3M-february-2018.txt’
~~~

The barcodes were then translated (8th and 9th base were changed to their complementary bases) to match the barcode cells of the Gene Expression part. The reads containing the expression part were processed with STARSolo version 2.7.10b [36] using:

~~~
--sjdbOverhang 100
--sjdbGTFfile ‘input.gtf’
--soloType CB_UMI_Simple --soloCBwhitelist
‘cellranger_barcodes_3M-february-2018.txt’
--soloUMIlen 12 --soloUMIdedup 1MM_CR
--soloUMIfiltering -
--soloCellFilter None
--outSAMmapqUnique 60
~~~

The same GTF file [37] used for BRB-seq was applied. Barcodes associated with empty droplets were filtered with `DropletUtils` [42] using the `EmptyDrops` method with a lower-bound threshold of 100 and a false discovery rate (FDR) threshold of 0.01.

Matrices were then processed with `Seurat` [43] version 4.3.0 in `R` version 4.3.0, following the methods described in Mayran et al. (2023). Barcodes with fewer than 200 identified genes and genes detected in fewer than three cells were filtered out. For CMO libraries, demultiplexing was performed in `R` using counts from `CITE-seq-Counts`. Cell barcodes with fewer than 5 CMO UMIs or absent in the `Seurat` object were discarded.

Sample attribution was performed using `demuxmix` [44] with the total number of UMIs per cell. Cells classified as non-singlets (negative, unsure, or doublets) were excluded. Low-quality cells and potential doublets were removed based on the mean UMI content and mitochondrial percentage. Barcodes with fewer than 0.4 times or more than 2.5 times the mean UMI, and those outside of 0.05% to 8% mitochondrial UMIs, were excluded.

The matrices were normalized, and the cell cycle score (using the 2019 updated gene list from `Seurat`) was computed. Samples were merged using the `merge` command in `Seurat`. The combined object was normalized, 2000 variable features were identified, and the data was scaled and regressed by cell cycle score and mitochondrial percentage. Principal components were computed using variable genes within the 5th and 80th percentiles of expression to limit batch effects. UMAP and k-nearest neighbors were computed with 25 principal components, and the clustering resolution (0.6) was optimized to avoid duplicate or missing clusters.

Cluster annotation was performed manually using marker genes. Genes from the module analysis of the BRB-seq experiment were scaled across the dataset and split to display each seeding number in Fig. 4E. The list of genes within each module was scored using the `addModuleScore` command in `Seurat`, and a custom `featurePlot` visualization was used as described in [13].

All NGS scripts generated for this study are deposited on GitHub: https://github.com/lldelisle/AllNGSscriptsFromBennabiEtAl2024.

## CODE ACCESSIBILITY

All the code required to reproduced the next generation sequencing analysis (bulk RNA-seq and scRNA-seq) can be found on: https://github.com/lldelisle/AllNGSscriptsFromBennabiEtAl2024 https://doi.org/10.5281/zenodo.14526130).

## Acknowledgements

We thank M.Cerminara, L. Friedman, A. Le Nabec, M. Nikolic, A.Shoushtarizadeh, and B.Zoller for their insightful comments and suggestions. This work was supported by Institut Pasteur (particularly the HPC core facility), Centre National de la Recherche Scientifique, CFM Foundation for Research, the French National Research Agency (ANR-20-CE12-0028’ChroDynE’ and ANR-23-CE13-0021’GastruCyp’ and ANR-10 LABX-73’Revive’), and by funding from the European Research Council (ERC-2023-SyG, Dynatrans, 101118866). I.B. was a recipient of a Revive postdoc fellowship. This work was also supported by the Ecole Polytechnique Fédérale de Lausanne and the Swiss National Science Foundation (SNSF, grant 310030 196868 to D.D., and grant 407940 206405 to A.M.) and the Human Frontier Science Program (HFSP LT000032/2019-L to A.M.). We thank the Gene Expression Research Core Facility (GECF) at EPFL for their expertise and service for sequencing experiments. The authors declare they have no competing interests.

## Author Contributions

I.B., P.H., A.M., and M.M. conceived the study and designed the experiments. I.B., P.H., A.M., and M.M. performed the experiments. D.K. and A.M. generated the mutant ESC lines. I.B., L.L.D., P.H., J.P., and A.M. analyzed the data. I.B., P.H., A.M., and T.G. wrote the manuscript. D.D. A.M. and T.G. secured funding and supervised the work.

## Notes

### Competing Interest Statement

The authors have declared no competing interest.

