## Supplementary Information for "Size-dependent temporal decoupling of morphogenesis and transcriptional programs in gastruloids"

**This PDF includes (in order of appearance):**
Supplementary Tables 1-6
Supplementary Figure 1-5

Table 1: Elongation timing statistics

| Sample | Replicate 1 | | Replicate 2 | | Replicate 3 | | Mesp2 reporter line | |
|---|---|---|---|---|---|---|---|---|
| | N | Elong. (h) | N | Elong. (h) | N | Elong. (h) | N | Elong. (h) |
| $N_0 = \mathbf{100}$ | 41 | 97 | 45 | 97 | 51 | n.a. | 41 | 101 |
| $N_0 = \mathbf{300}$ | 41 | 104 | 44 | 102 | 38 | 99 | 49 | 106 |
| $N_0 = \mathbf{600}$ | 36 | 110 | 43 | 111 | 27 | 102 | 48 | 114 |
| $N_0 = \mathbf{1200}$ | 40 | 119 | 39 | 114 | 32 | 106 | 52 | 116 |

Table 2: Uniaxiality timing statistics.

| Sample | Mean t (h) | S.d. (h) | C.v. (%) | Fraction uniaxial | Fraction uniaxial (%) |
|---|---|---|---|---|---|
| $N_0 = \mathbf{150}$ | 103.02 | 1.30 | 1.26 | 41 / 41 | 100.00 |
| $N_0 = \mathbf{300}$ | 108.04 | 6.71 | 6.21 | 49 / 49 | 100.00 |
| $N_0 = \mathbf{600}$ | 115.04 | 9.17 | 7.97 | 47 / 48 | 97.92 |
| $N_0 = \mathbf{1200}$ | 125.84 | 7.77 | 6.17 | 50 / 52 | 96.15 |

Table 3: Polarization timing statistics.

| Sample | Mean t (h) | S.d. (h) | C.v. (%) | N |
|---|---|---|---|---|
| $N_0$ = **150** | 103.02 | 1.30 | 1.26 | 41 |
| $N_0$ = **300** | 104.78 | 1.31 | 1.25 | 49 |
| $N_0$ = **600** | 105.29 | 1.54 | 1.46 | 48 |
| $N_0$ = **1200** | 106.40 | 1.96 | 1.85 | 52 |

Table 4: Primers used for genotyping Mesp locus and GFP/mCherry integrations.

| Primer | Sequence | Expected size | Function |
|---|---|---|---|
| Fw_aroundDel | AAGCCTTCCTCTGGATAC | 784 | Genotyping Mesp locus deletion |
| Rv_aroundDel | CCTGTGACCGTAGGTTTA | | |
| Fw_Ar._GFP | GCCTCTTCCAGGATCTAC | 1174 | Genotyping Mesp1 GFP integration |
| Rv_Ar._GFP | GACATGCTGGCTCTTCTA | | |
| Fw_Ar._M1-GFP | AAGCCTTACATGGCTCAG | 2884 | Genotyping Mesp1 GFP integration |
| Rv_Ar._M1-GFP | CGGCTTTGATATGCTGATG | | |
| Fw_Ar._M1-5' | CCTTGCTCACCATGATATAAG | 1146 | Genotyping Mesp1 GFP integration |
| Rv_Ar._M1-5' | CCTAGGGCTCAGGATAAAG | | |
| Fw_Ar._mCherry | TGGCTGTCCTGAACTTTG | 1135 | Genotyping Mesp2 mCherry integration |
| Rv_Ar._mCherry | AGGCAGATAAAGGCACTTC | | |
| Fw_Ar._M2-5 | TGGCTGTCCTGAACTTTG | 1829 | Genotyping Mesp2 mCherry integration |
| Rv_Ar._M2-5 | CCTTGGCAGGCTTAGATT | | |

Table 5: List of single guide RNAs

| single guide function | 5'sgRNA | 3'sgRNA |
|---|---|---|
| Mesp_deletion | GAGGCAGCCGCCTGTCTTCG | AAGCGGGGACCCTCCTGTGG |
| Mesp1_GFP_Targeting | GAGTGGAGGGGACAATGCAA | |
| Mesp2_mCherry_Targeting | GCAGCCCAAATCCACACCGCA | |

Table 6: Primary and secondary antibodies and fluorescent dye specifications.

| Antibody | Species | Reference | Provider | Dilution |
|---|---|---|---|---|
| BRA | Goat | AF2085 | RnD Systems | 1:100 |
| FOXC1 | Rabbit | EPR20678 | Abcam | 1:500 |
| SOX2 | Rat | 14-9811-82 | eBioscience | 1:200 |
| anti-goat AF-488 | Donkey | A-11055 | Invitrogen | 1:500 |
| anti-rabbit AF-546 | Donkey | A-11010 | Invitrogen | 1:500 |
| anti-rat AF-647 | Donkey | A78947 | Invitrogen | 1:500 |
| DAPI | - | D3571 | Invitrogen | 1:500 |

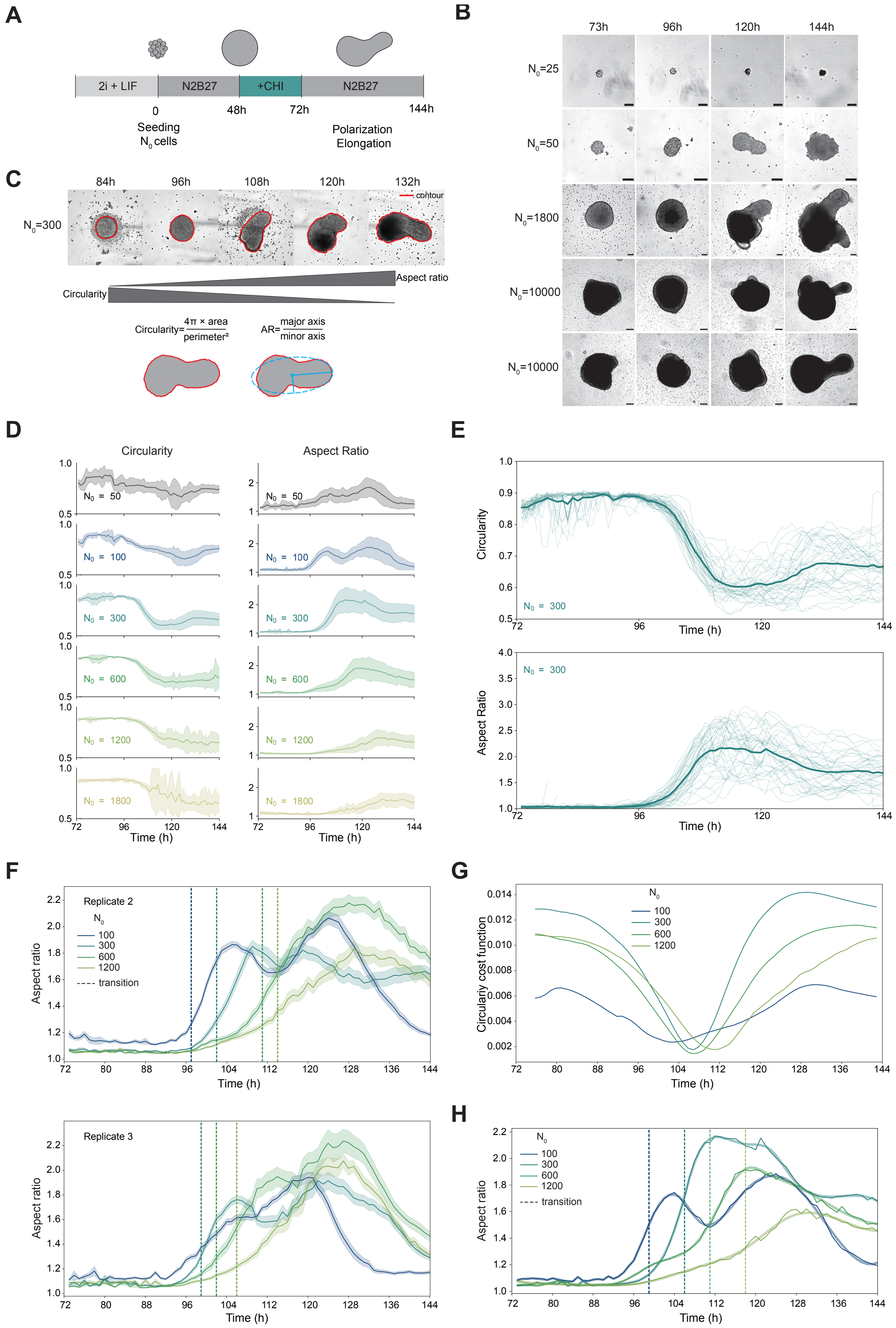


**See next page for caption of Fig. S1.**

**Supplementary Figure 1**: **Experimental protocol and morphological quantification.** (A) Gastruloid protocol: Initial cell numbers ($N_0$) of mESCs were seeded and subjected to a WNT activation pulse (Chiron) from 48 to 72 h after seeding. Gastruloids progressively break symmetry and elongate an anterior-posterior axis. (B) Brightfield images illustrating size limits for gastruloid growth and elongation. Scale bars = 100 $\mu$m. (C) Automated analysis of time-lapse images detects gastruloid contours, with shape described by circularity (1 = perfect circle, 0 = elongated) and aspect ratio (1 = perfect circle, >> 1 = elongated). (D) Temporal dynamics of circularity and aspect ratio (mean $\pm$ s.d.) across $N_0$ conditions. Dashed lines indicate transition times. Data are from a single representative experiment. Colors represent $N_0$ = 50 (n = 44), $N_0$ = 100 (n = 41), $N_0$ = 300 (n = 41), $N_0$ = 600 (n = 36), $N_0$ = 1200 (n = 40), and $N_0$ = 1800 (n = 25). (E) Individual time course of circularity (top) and aspect ratio (bottom) for gastruloids seeded with $N_0$ = 300 cells, tracked from 72 h to 144 h post-seeding. Each line represents a gastruloid; bold line is the mean. (F) Mean shape descriptors (mean $\pm$ s.e.m.) across time (post-seeding) for varying $N_0$. Data are for two additional experimental replicates. See Supplementary Table 1 for sample numbers. (G) Circularity cost function minima marking transition times for symmetry breaking (Methods). (H) Aspect ratio spline fitting identifies transition times (dashed lines), consistent with times evaluated using optimal partitioning (Fig. 1F). Elongation transitions occur at 99 h, 106 h, 111 h, and 118 h for $N_0$ = 100, 300, 600, and 1200, respectively.

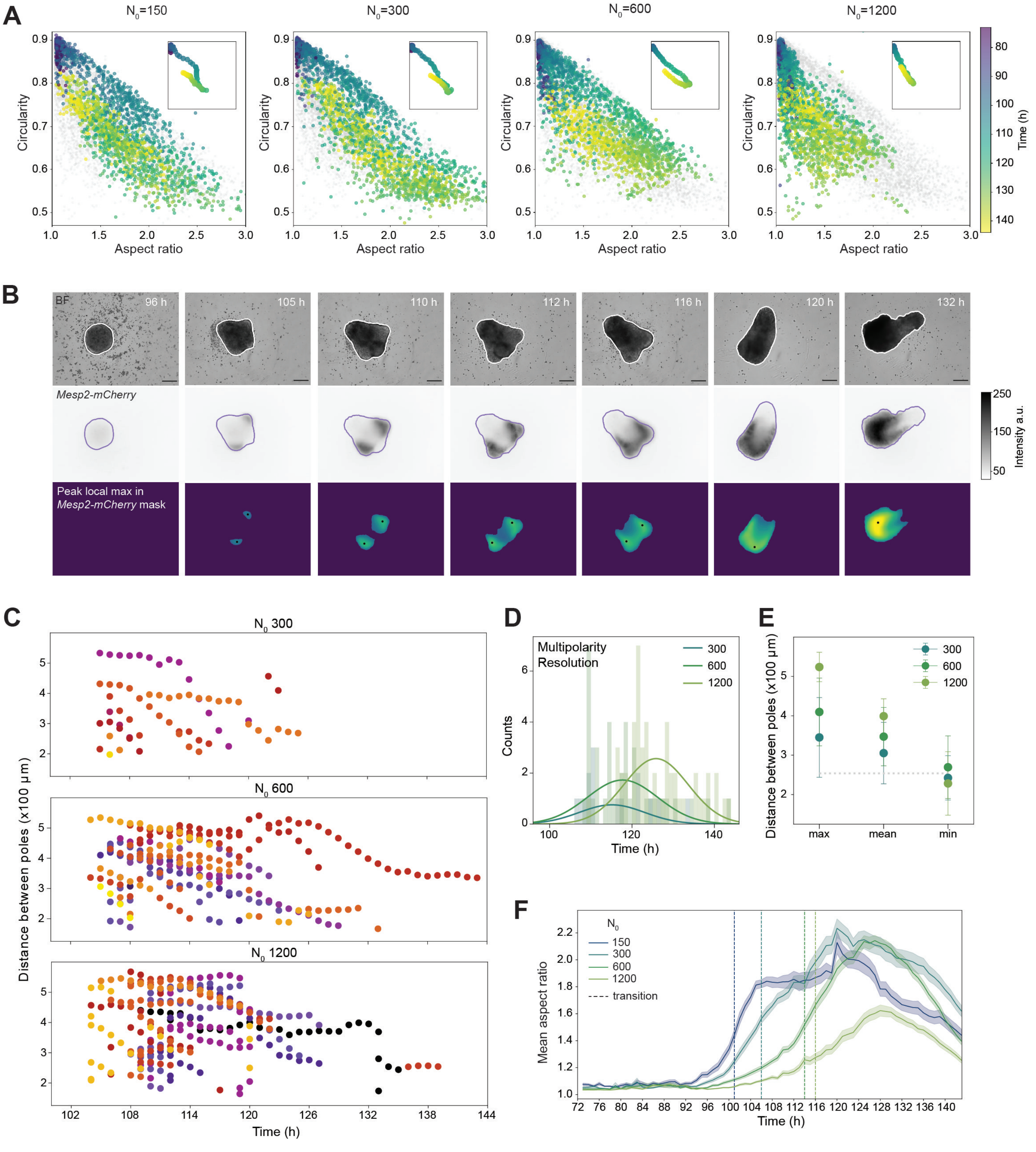


**See next page for caption of Fig. S2.**

**Supplementary Figure 2**: **Quantification of multipolarity and *Mesp2* expression dynamics.** (A) Scatter plots of circularity vs. aspect ratio for gastruloids with varying $N_0$, colored by time (72–144 h). Insets highlight average morphological trajectories. Data are from one representative experiment with $N_0 = 100$ (n = 41), $N_0 = 300$ (n = 49), $N_0 = 600$ (n = 48), and $N_0 = 1200$ (n = 52). (B,C) Temporal dynamics of circularity and aspect ratio (mean ± s.e.m.) across $N_0$ conditions. Dashed lines mark symmetry breaking and elongation transitions, determined via optimal partitioning (Fig. S1G and methods). Symmetry breaking (from circularity) occurs at 103 h, 110 h, 115 h, and 115 h for $N_0 = 100$, 300, 600, and 1200, respectively; elongation (from aspect ratio) occurs at 101 h, 106 h, 114 h, and 116 h. (D) Example analysis of multipolarity in $N_0 = 600$ gastruloids over time. Brightfield images (top), *Mesp2*-mCherry expression (middle), and *Mesp2*-mCherry-positive area and local maxima analysis (bottom) are shown. Whole gastruloid contour extracted from top (white line) overlaid in middle (purple line). Scale bars = 200 $\mu$m. (E) Euclidean distances between peak local maxima over time for multipolar gastruloids from different $N_0$. Colors indicate individually followed gastruloids. (F) Histograms of multipolarity resolution times (transition to a single *Mesp2* pole) for each $N_0$; solid lines show Gaussian fits (see Supplementary Tables for statistics). (G) Maximum, mean, and minimum distances between *Mesp2* poles for each $N_0$. Gray dashed line is the average minimum distance across all gastruloids, min = 253 ± 77 $\mu$m (mean ± s.d.).

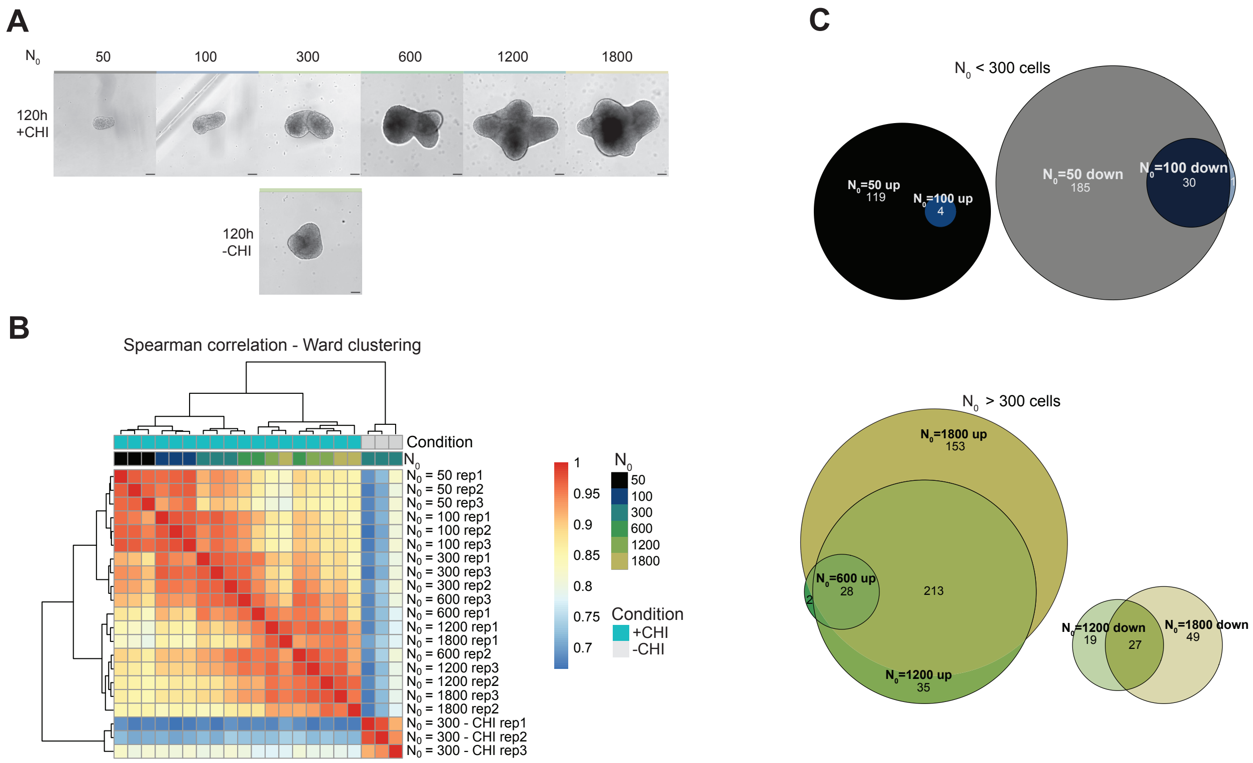


**Supplementary Figure 3**: **Transcriptional profiling of gastruloids across sizes.** (A) Brightfield images of gastruloids at 120 h across varying $N_0$, taken from one experimental batch used for bulk RNA-seq (Fig.3). Scale bars = 100 $\mu m$. (B) Correlation matrix of samples clustered by the 1000 most variable genes. Colors are Spearman correlation values; clustering uses the Ward D2 algorithm. (C) Venn diagrams of misregulated genes across size comparisons: gastruloids $<$ 300 cells (top), gastruloids $>$ 300 cells (bottom).

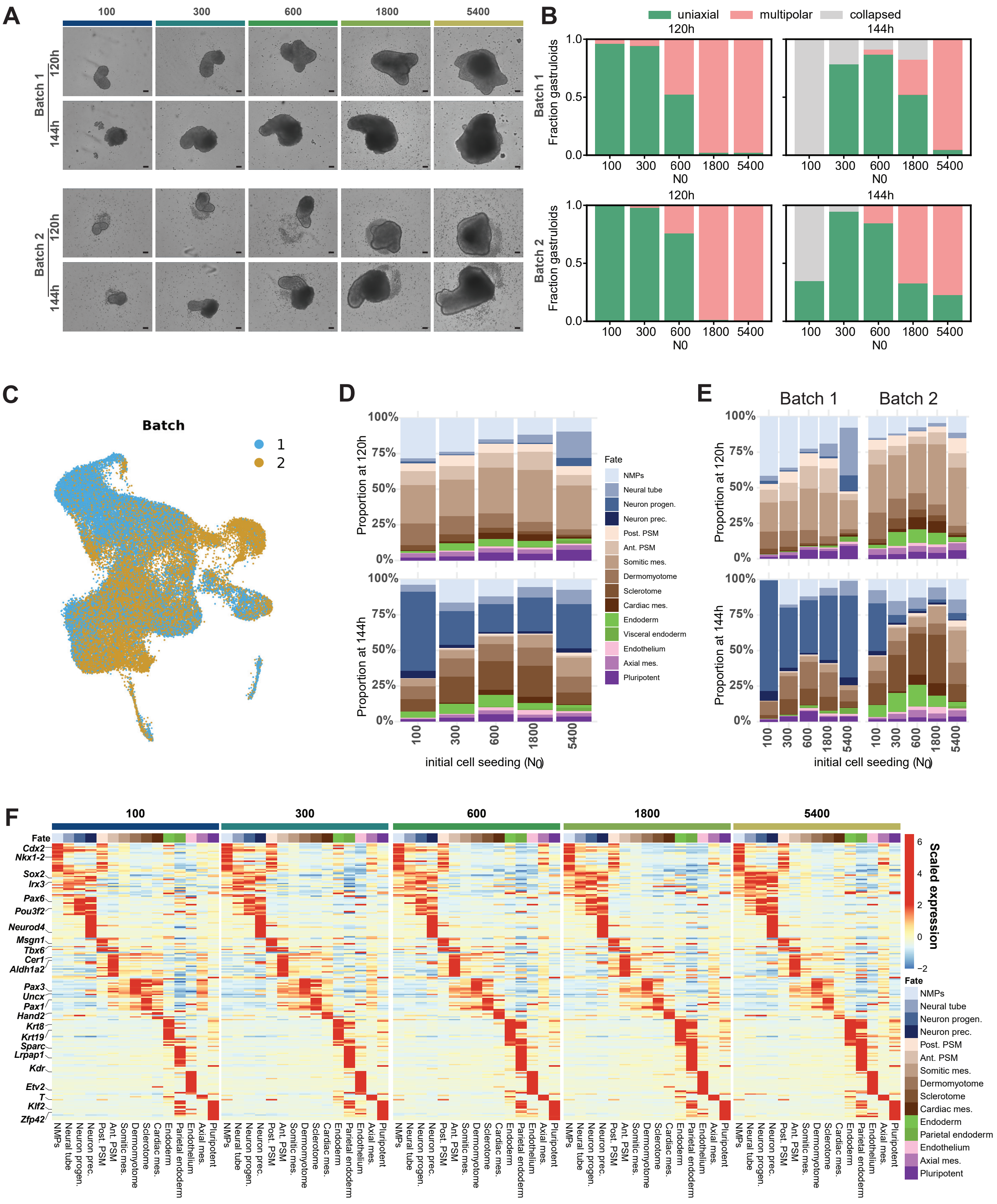


**Supplementary Figure 4: Batch-batch differences exceed size-dependent variations in cell fate composition.** (A) Brightfield images of gastruloids from indicated $N_0$ across two experimental batches used in the single-cell analysis in Fig. 4. Same gastruloids shown at 120 h (top) and 144 h (bottom). Scale bar = 50 $\mu m$. (B) Proportion of gastruloids manually categorized as uniaxial, multipolar, or collapsed (for two experimental batches). (C) UMAP projections of data shown in Fig. 4, colored by experimental batch. (D-E) Bar plots of cell identity proportions from single-cell analysis split by time after aggregation and $N_0$. Two batches are merged together (D) or separated (E). (F) Heatmap of pseudo-bulk expression for marker genes of the identified clusters. Data is scaled across the entire dataset, with separate panels for each $N_0$.

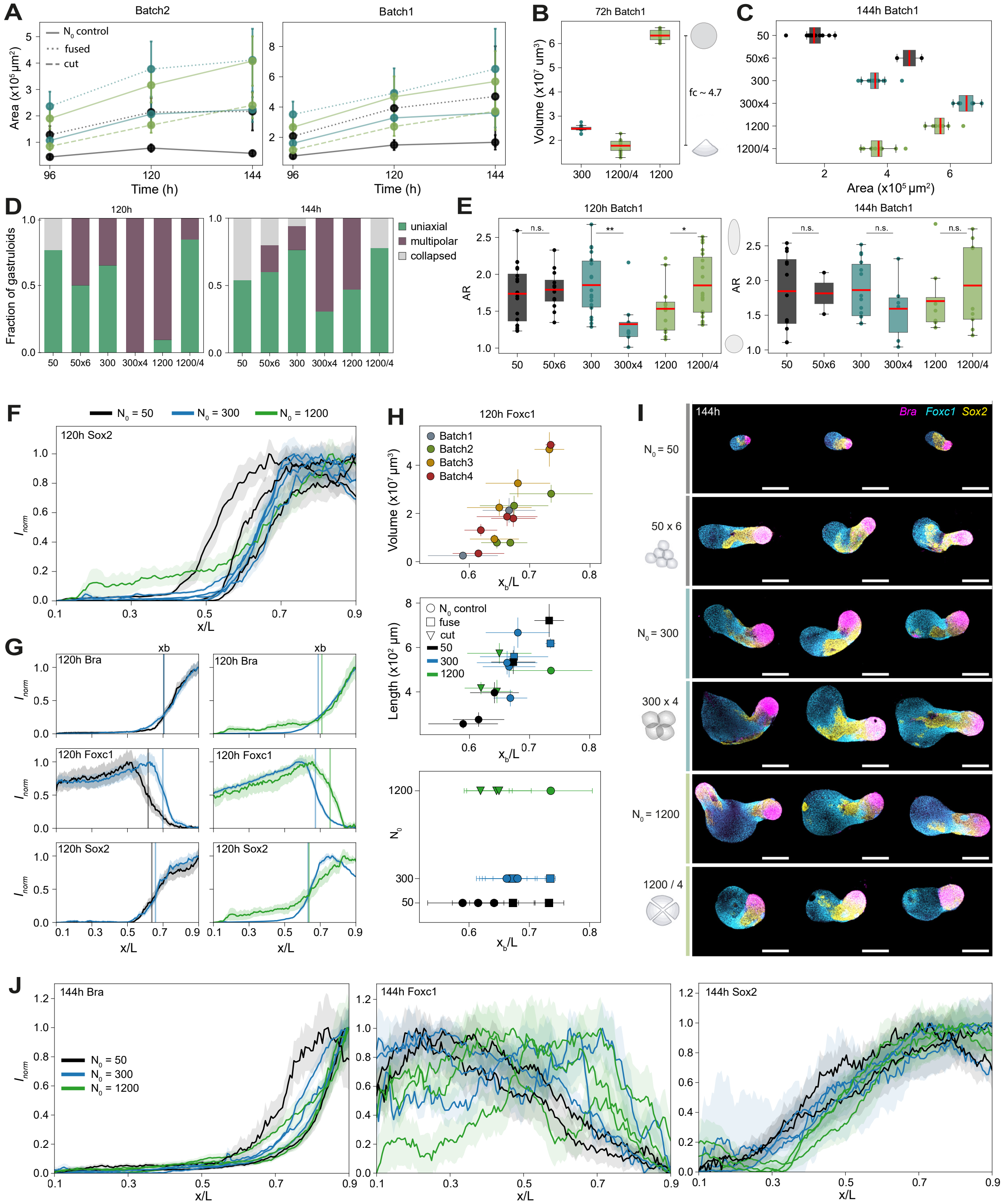


**See next page for caption of Fig. S5.**

**Supplementary Figure 5**: **Expanded effects of gastruloid resizing on morphogenesis and patterning.** (A) Mean gastruloid area as 96 h, 120 h, and 144 h post-manipulation across two experimental batches. Colors correspond to $N_0$; error bars are s.d. (B) Mean gastruloid area at 72 h immediately after perturbation for $N_0 = 300$, $N_0 = 1200$, and cut $N_0 = 1200$ (1200/4). Colors correspond to $N_0$; error bars are s.d. (C) Boxplot of gastruloid area at 144 h (as in Fig. 5D) for the second experiment batch. (D) Proportions of manually annotated gastruloids categorized as uniaxial, multipolar, or collapsed at 120 h and 144 h for the second experiment batch. (E) Gastruloid aspect ratio (proxy for uniaxial elongation) at 120 h and 144 h (as in Fig. 5F). P-values from a two-sided t-test indicated: <0.05 (*), <0.01 (**), and <0.001 (***), n.s. = not significant. (F) Normalized mean expression profiles (mean ± s.e.m.) of Sox2 at 120 h as a function of the relative position (x/L) as described in Fig. 5H; results of four experimental batches; AP-axis oriented left-to-right. (G) Mean expression profiles of Bra, Foxc1, and Sox2 at 120 h (as in Fig. 5H), showing variability between conditions within a single experiment (left: batch 4, right: batch 2). Vertical lines indicate pattern boundary positions (xb) of the mean profiles marked at the half-maximal expression value (EC50). (H) Scatter plots showing the relationship between the mean pattern boundary positions (xb) of Foxc1 and gastruloid volume (top), length (middle), and $N_0$ (bottom). Error bars are s.d. on both axes. (I) Maximum projections of confocal image stacks of 144 h gastruloids immunofluorescently stained for Bra, Foxc1, and Sox2. Posterior ends oriented to the right. Scale bars = 200 $\mu m$. (J) Normalized mean expression profiles (mean ± s.e.m.) of Bra, Foxc1, and Sox2 at 144 h as a function of the relative position (x/L) (as in Fig. 5H). Results from four experimental batches are shown, with the AP-axis oriented left-to-right.
